# A fluorescent reporter for FtsA is functional as the sole FtsA in *Escherichia coli* and has hypermorphic properties

**DOI:** 10.1101/2022.10.31.514644

**Authors:** Todd A. Cameron, William Margolin

## Abstract

FtsA, a homolog of actin, is essential for cell division of *Escherichia coli* and is widely conserved among many bacteria. FtsA helps to tether polymers of the bacterial tubulin homolog FtsZ to the cytoplasmic membrane as part of the cytokinetic Z ring. GFP fusions to FtsA have illuminated FtsA’s localization in live *E. coli*, but these fusions have not been fully functional and required the presence of the native FtsA. Here, we characterize “sandwich” fusions of *E. coli* FtsA to either mCherry or msfGFP that are fully functional for cell division and exhibit fluorescent rings at midcell that persist throughout constriction until cell separation. FtsA within the Z ring moved circumferentially like FtsZ, and FtsA outside the rings formed highly dynamic patches at the membrane. Notably, both FtsA-mCherry and FtsA-msfGFP acted as mild hypermorphs, as they were not toxic when overproduced, bypassed the essential cell division protein ZipA, and suppressed several thermosensitive *fts* alleles, although not as effectively as the prototypical hypermorph FtsA*. Overall, our results indicate that fluorescent FtsA sandwich fusions can be used as the sole FtsA in *E. coli* and thus should shed new light on FtsA dynamics during the cell division cycle in this model system.

**Importance:** FtsA is a key conserved cell division protein, and *E. coli* is the most well studied model system for bacterial cell division. One obstacle to full understanding of this process is the lack of a fully functional fluorescent reporter for FtsA *in vivo*. Here, we describe a fluorescent fusion to *E. coli* FtsA that divides cells efficiently in the absence of the native FtsA and can be used to monitor FtsA dynamics during cell division.

## INTRODUCTION

To divide one cell into two daughter cells, bacteria must interleave the biosynthetic and regulatory activities of dozens of cytoplasmic and periplasmic proteins into a singular, organized divisome. The bacterial tubulin homolog FtsZ plays a central role in establishing and maintaining this divisome by forming an “Z ring” consisting of multiple treadmilling FtsZ filaments that move along the cytoplasmic membrane in line with the future plane of division (Moore *et al*., 2017; Yang *et al*., 2017). By repeatedly interacting with other divisome proteins as they travel, these FtsZ filaments establish the fundamental organization for the entire divisome (Baranova *et al*., 2020).

FtsZ lacks intrinsic affinity for the membrane surface and must rely on interactions with its membrane anchors for cells to progress through cell division (Pichoff and Lutkenhaus, 2002). Interactions with FtsA, the primary such anchor, are mediated by the conserved C-terminal peptide of FtsZ (Din *et al*., 1998; Ma and Margolin, 1999). FtsA in turn remains closely associated with the membrane due to its amphipathic C-terminal tail (Pichoff and Lutkenhaus, 2005). In *E. coli* and other Gammaproteobacteria, FtsZ also interacts with an additional essential membrane anchor ZipA (Pichoff and Lutkenhaus, 2002). Although both proteins serve as FtsZ membrane anchors, FtsA has additional critical roles recruiting or interacting with downstream divisome proteins such as FtsN, FtsK, FtsEX, and FtsW (Wang and Lutkenhaus, 1998; Bernard *et al*., 2007; Du *et al*., 2019; Park *et al*., 2021),

As an actin homolog, FtsA is also capable of forming ATP-dependent polymeric structures. However, since FtsA is far less abundant than FtsZ, with ∼1000 FtsA proteins per *E. coli* cell compared to ∼7000 of FtsZ (Li *et al*., 2014), FtsA is most likely distributed across FtsZ filaments as individual proteins or small oligomers rather than as long filaments. There is growing evidence that *E. coli* FtsA can form several different polymeric structures, including minirings and antiparallel filaments, that serve distinct structural and regulatory roles throughout cell division.

As shown on lipid monolayers, wild-type (WT) *E. coli* FtsA forms dodecameric minirings that loosely separate FtsZ filaments (Krupka *et al*., 2017). In contrast, hypermorphic FtsA* mutants, which can bypass the absence of the otherwise essential ZipA or FtsEX proteins *in vivo*, instead form more highly packed arcs or double filaments on lipids *in vitro* and promote FtsZ filament bundling (Krupka *et al*., 2017; Schoenemann *et al*., 2018). It has therefore been hypothesized that FtsA minirings might be disrupted through a ZipA- or FtsEX-dependent mechanism (Du *et al*., 2016; Vega and Margolin, 2019), thus allowing minirings to function as early divisome checkpoints by limiting FtsZ bundling, by occluding sites on FtsA that recruit downstream divisome proteins, or both.

Subsequent interactions between FtsA and one such divisome protein, FtsN, seems to enhance assembly of FtsA double filament structures. The short cytoplasmic tail of FtsN was recently found to be sufficient to induce FtsA to form short, antiparallel filaments on lipid monolayers (Nierhaus *et al*., 2022). These filaments prefer regions of negative membrane curvature and may therefore help FtsA to align FtsZ filaments circumferentially around the invaginating division septum (Nierhaus *et al*., 2022). Such a function would closely resemble a similar role served by another actin-like protein, MreB, for cell elongation.

Fluorescent protein fusions to divisome components have provided critical insights into the process of cell division, allowing for protein behaviors to be monitored in real time and permitting even the movement of individual protein molecules to be tracked. Unfortunately, as observed starting with the initial FtsA-GFP fusion made in the mid 1990’s and in subsequent studies, fluorescent protein fusions to either the FtsA C-terminus or N-terminus are not fully functional and fail to complement the loss of native *ftsA* (Ma *et al*., 1996; Pichoff and Lutkenhaus, 2005). Given the multiple lateral and longitudinal FtsA-FtsA contacts required for polymeric FtsA structures, as well as its interactions with other cell division proteins, FtsA is likely particularly sensitive to the inopportune placement of a bulky GFP domain. Although these fusions do localize to the Z ring, the lack of complementation makes it unclear whether they accurately reflect the activity and localization of native FtsA.

In cases such as this, an alternate ‘sandwich fusion’ approach has sometimes proven useful creating functioning fluorescent fusions. Rather than creating N- or C-terminal fusions, this strategy targets surface-exposed loops for an internal insertion of the fluorophore sequence, thus allowing for the bulky fluorescent protein to be repositioned to potentially less disruptive locations around the target protein. This approach was previously employed to generate functional, fluorescently tagged constructs of *E. coli* MreB and FtsZ (Bendezú *et al*., 2009; Moore *et al*., 2017) as well as *Bacillus subtilis* FtsA (Bisson-Filho *et al*., 2017).

Here we report the successful creation of two such sandwich fusions between *E. coli* FtsA and either mCherry or monomeric superfolder GFP (msfGFP). Unlike previous fusions to *E. coli* FtsA, these can complement an FtsA null mutation and maintain WT cell morphology, although each acts as a weak FtsA hypermorph and can suppress several divisome defects. Using these fusions, we found that FtsA within the Z ring moves circumferentially at a rate similar to FtsZ whereas FtsA outside the rings forms highly dynamic patches at the membrane. Overall, our results demonstrate that these fluorescent FtsA sandwich fusions are functional for cell division in *E. coli* and can be used to shed new light on FtsA dynamics during the cell division cycle in this model system.

## RESULTS and DISCUSSION

### Fluorescent protein insertions in FtsA are functional in the absence of native FtsA

As MreB and *B. subtilis* FtsA remained functional with insertions of monomeric fluorescent proteins in the surface-exposed loop between helix 7 and sheet 12 in domain 2B, we chose to insert msfGFP and mCherry between S267 and I268 within this region of *E. coli* FtsA (Fig. 1A-B). In keeping with the nomenclature of previously reported “sandwich” fusions, we named these FtsA-msfGFP^sw^ and FtsA-mCherry^sw^, respectively. We also included short peptide linkers flanking the fluorescent protein insertion to allow additional flexibility (Fig. 1A).

**Fig. 1.**
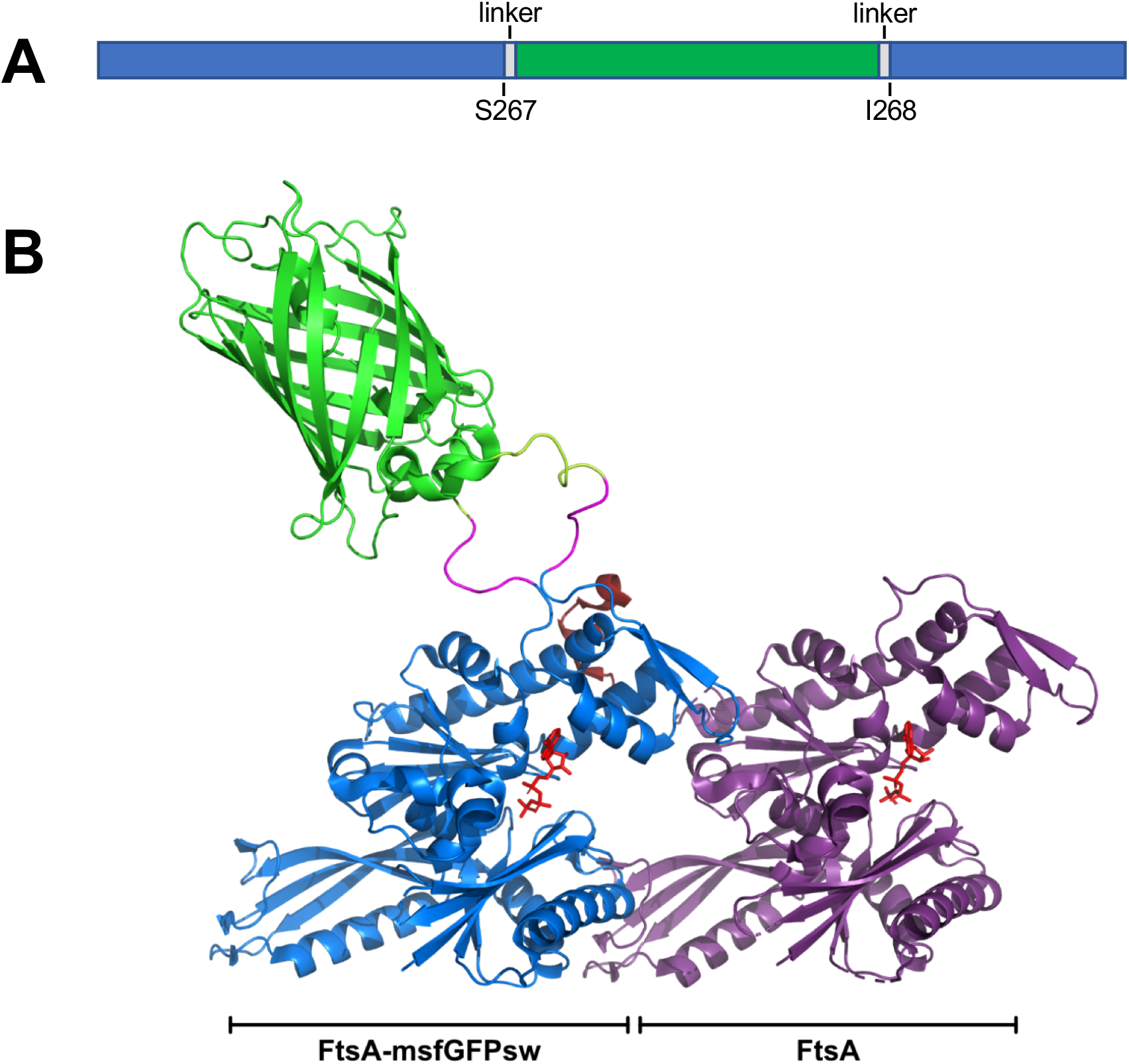
Engineering fluorescent protein sandwich fusions to *E. coli* FtsA. A. Schematic of msfGFP^sw^ insertion in the ftsA gene, showing encoded residues. B. Structural model for insertion of msfGFP or mCherry into FtsA, using the *Thermotoga maritima* structure (1E4G).

Expression of either fusion protein from the IPTG-inducible weakened *trc* promoter on the pDSW210 plasmid in WT *E. coli* cells resulted in strong localization to the divisome similar to previous GFP fusions to the termini of FtsA, indicating that the sandwich fusion did not interfere with the ability of FtsA to interact with FtsZ and the Z ring (see below). We then tested whether the sandwich fusions were functional, either by complementing a thermosensitive *ftsA12* mutant with the plasmid or introducing a frameshift mutation in the chromosomal *ftsA* gene (*ftsA*^*o*^) by transduction with a linked Tn*10* (tetracycline resistance) marker. Although WT FtsA and the hypermorphic FtsA* (R286W) mutant expressed from pDSW210 could complement *ftsA12* at the nonpermissive temperature of 42°C without any IPTG induction, the mCherry^sw^ and msfGFP^sw^ fusions could not complement unless at least 50 μM IPTG was added (Fig. 2A, middle rows), indicating that these sandwich fusions may be partially thermosensitive. Notably, 50 μM IPTG expresses sufficient WT FtsA to inhibit viability, consistent with the need to keep the FtsZ:FtsA ratio within a normal range (Dai and Lutkenhaus, 1992; Dewar *et al*., 1992). In contrast, there were no growth defects when either sandwich fusion or FtsA* was grown with 50 μM or even 200 μM IPTG.

**Fig. 2.**
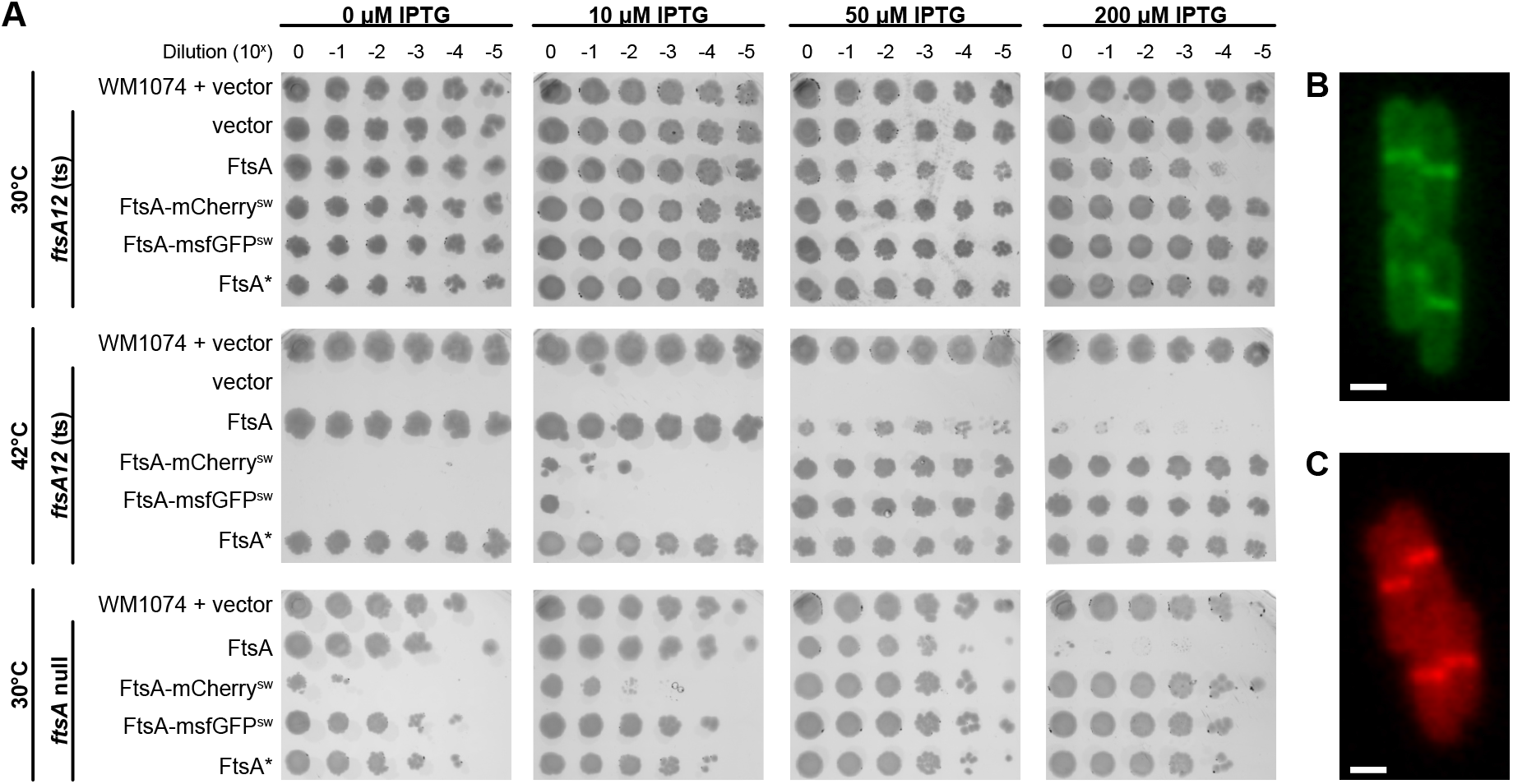
Sandwich fusions of FtsA to msfGFP or mCherry are functional and form sharp midcell bands in cells lacking native FtsA. (A) Viability of WM1074 containing native *ftsA* and pDSW210 compared to thermosensitive *ftsA12* strains and *ftsA*^*0*^ strains containing pDSW210, pDSW210-FtsA, pDSW210-FtsA*, pDSW210-FtsA-mCherry^sw^, or pDSW210-FtsA-msfGFP^sw^. Serial dilutions were spotted on plates containing various IPTG concentrations (top). *ftsA12* strains were grown at both permissive and non-permissive temperatures. Note that WT FtsA starts to inhibit cell division when expressed at higher levels. (B) Fluorescence micrograph of WM6246 (*ftsA*^*0*^ pDSW210-FtsA-msfGFP^sw^) cells grown to mid-logarithmic phase the presence of 50 μM IPTG showing localization of FtsA-msfGFP^sw^ to midcell rings in dividing cells. (C) Fluorescence micrograph of WM4601 (*ftsA*^*0*^ pDSW210-FtsA-mCherry^sw^) cells grown to mid-logarithmic phase in the presence of 50 μM IPTG showing localization of FtsA-mCherry^sw^ to midcell rings. Scale bar, 1 μm.

In keeping with the idea that the fusion proteins might be partially thermosensitive but otherwise functional, cells expressing FtsA-msfGFP^sw^ or FtsA-mCherry^sw^ could be transduced with the *ftsA*^*o*^ allele at normal efficiencies (∼50% linkage with a *leuO*::Tn*10* marker) in the presence of IPTG (data not shown). However, whereas FtsA-msfGFP^sw^ was fully viable as the sole source of FtsA in the absence of IPTG at 30°C, FtsA-mCherry^sw^ required some IPTG induction for full viability in the *ftsA*^*o*^ background (Fig. 2A, bottom rows). Cells with the *ftsA*^*o*^ allele that expressed either sandwich fusion in the presence of IPTG were short, with no sign of cell division inhibition, and had sharply localized fluorescent rings at midcell (Fig. 2B-C). This indicates that the sandwich fusions are functional as the sole copy of FtsA in the cell and contrasts with either lack of full functionality or dominant negative effects of N or C-terminal fluorescent protein fusions to FtsA when expressed from plasmids (Ma *et al*., 1996; Pichoff and Lutkenhaus, 2005).

### Further exploration of fusion protein functionality

Although the sandwich fusions seemed to be somewhat thermosensitive, their ability to fully complement the *ftsA*^*o*^ allele, particularly the msfGFP^sw^ fusion without IPTG, indicated that they were potentially as functional as WT FtsA. Immunoblot analysis indicated that uninduced levels of WT FtsA produced from pDSW210 were ∼4 fold higher than native levels, consistent with the ability of WT FtsA to complement without IPTG induction (Fig. 3). Similarly, uninduced levels of the msfGFP and mCherry fusions were 3-4-fold higher than native levels. The inability of FtsA-mCherry^sw^ to complement at these levels unless IPTG was added suggests that this fusion is less active than FtsA-msfGFP^sw^ or WT FtsA. In comparison, levels of WT FtsA or the sandwich fusions induced at 100 μM IPTG were ∼10 fold higher than native levels (Fig. 3).

**Fig. 3.**
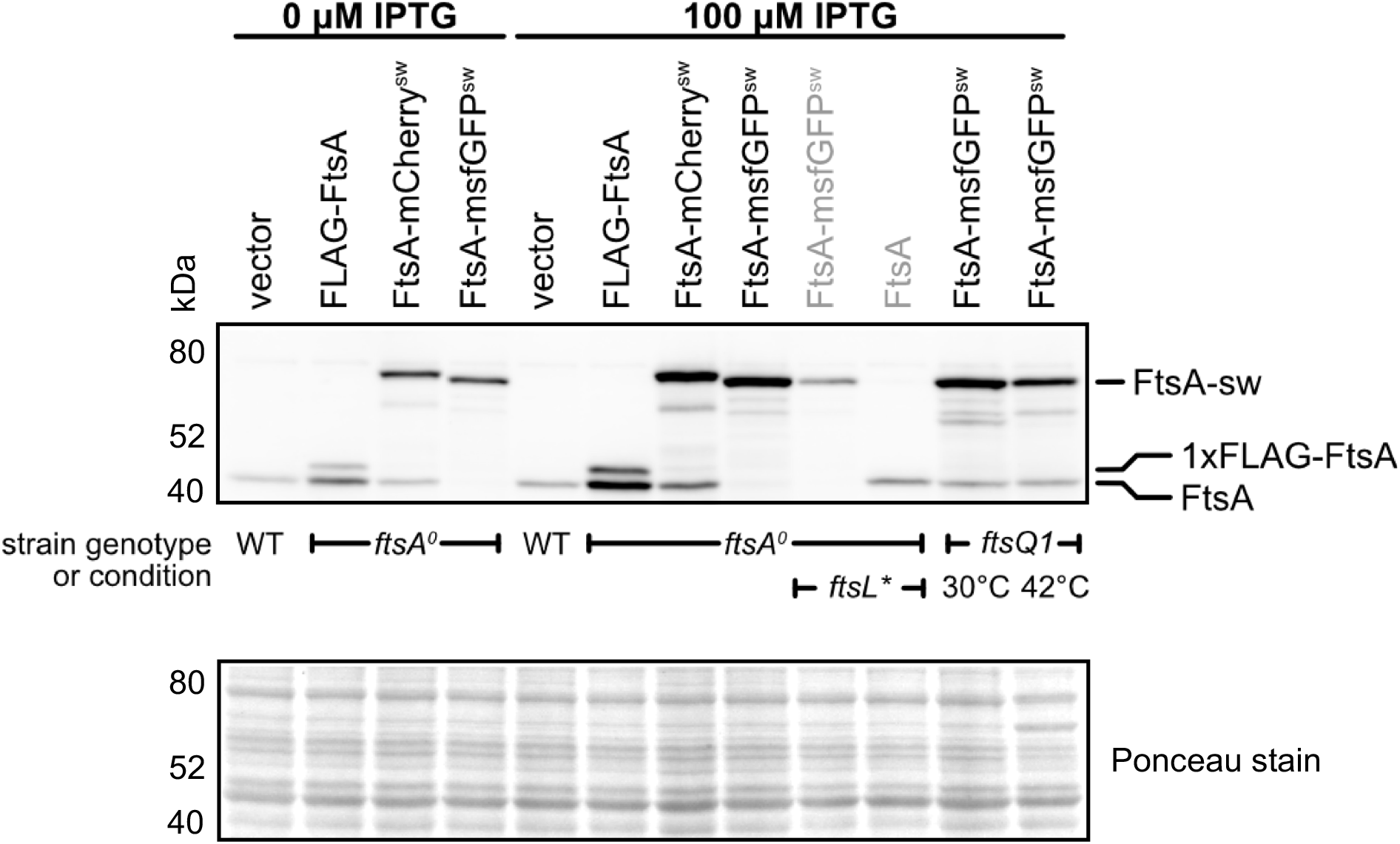
Expression of FtsA sandwich fusions under different conditions. Expression of FLAG-FtsA or FtsA sandwich fusions from plasmids in strains with genotypes depicted at bottom of the blot (top) were grown to mid-logarithmic phase ± 100 μM IPTG at 30°C (or 42°C for 1 h for the rightmost lane) and were probed with anti-FtsA antibody on immunoblots. All plasmids were pDSW210 (labeled with black letters) except for weaker-expressing pSEB440 derivatives (labeled with gray letters). Bands corresponding to FtsA sandwich fusions (FtsA-sw), 1XFLAG-FtsA and FtsA are highlighted. Shown at the bottom is the same blot stained with Ponceau as a loading/transfer control.

To explore further the correlation between expression and function and to define the minimum level of FtsA needed for normal cell division, we attempted to lower expression of WT *ftsA* from pDSW210 by further weakening the *trc* promoter, weakening the ribosome binding site (RBS), or engineering a stronger *lac* operator. However, in each case WT FtsA expressed from these altered plasmids could still complement the *ftsA*^*o*^ allele without IPTG induction (data not shown).

To circumvent these issues, we obtained a plasmid, pSEB440, from the Lutkenhaus laboratory that expresses FtsA at a much lower level than pDSW210 from an IPTG-inducible promoter on a lower-copy, p15a replicon. Unlike pDSW210-FtsA, pSEB440 cannot complement the *ftsA*^*o*^ allele unless induced with greater than ∼125 μM IPTG (Pichoff *et al*., 2018) (data not shown), making it a more sensitive system for measuring the lower limits of FtsA needed for function. We found that the likely reason for the low expression from pSEB440, in addition to its weakened *trc* promoter and RBS, is the existence of a small open reading frame (ORF), ATG GAA TAG, immediately downstream from the RBS. This is followed by a 16-bp linker, which leads to the ATG for the *ftsA* ORF. The presence of this short upstream ORF may decrease translation of FtsA sufficiently to dampen its expression.

We then replaced the WT FtsA in pSEB440 with FtsA-msfGFP^sw^ to directly compare its ability to complement with that of WT FtsA. We were surprised to find that *ftsA*^*o*^ could not be introduced into cells carrying pSEB440-FtsA-msfGFP^sw^, even with IPTG induction at 0.1 or 1 mM, suggesting either that the levels of fusion protein from this plasmid are lower than those of WT FtsA at a given IPTG induction level, or that FtsA-msfGFP^sw^ has lower specific activity than WT FtsA. Immunoblotting indicated that the second possibility was most likely, as the WT FtsA or FtsA-msfGFP^sw^ protein bands produced from pSEB440 induced with 0.1 mM IPTG had equivalent intensities (less than 2-fold higher than native FtsA levels) and were significantly lower in intensity than the same proteins produced from uninduced pDSW210 (Fig. 3), which were ∼ 4-fold higher than native levels, as mentioned previously.

These results therefore suggested that FtsA-msfGFP^sw^ has lower specific activity than WT FtsA. In support of this idea, we could only obtain *ftsA*^*o*^ transductants of cells expressing FtsA-msfGFP^sw^ from pSEB440 if the *ftsL*\* hyperfission allele (E88K) was co-transduced with *ftsA*^*o*^ in the presence of IPTG (data not shown). Previously it was found that levels of FtsA that were too low to complement a WT strain regained their ability to complement upon introduction of the *ftsL*\* allele (Tsang and Bernhardt, 2015; Park *et al*., 2020). Taken together, these results suggest that activation of the divisome by *ftsL*\* reduces the number of FtsA-msfGFP^sw^ molecules needed for function in the same way that the minimum requirement for WT FtsA levels is reduced.

### FtsA sandwich fusions have hypermorphic properties

It is well known that higher expression of WT FtsA inhibits cell division due to the importance of the FtsZ:FtsA ratio as mentioned above. However, we noticed that higher levels of IPTG induction of the pDSW210 plasmid derivatives, resulting in 10-fold or higher levels relative to native FtsA levels (Fig. 3) did not affect viability or division of cells expressing FtsA-mCherry^sw^ or FtsA-msfGFP^sw^ whereas cell division is strongly inhibited with the equivalent expression of WT FtsA (Fig. 2A and data not shown). We suspected that this lack of toxicity could stem from hypermorphic properties of these fusion proteins, as hypermorphic FtsA* and FtsA*-like derivatives are often not toxic when overproduced (Geissler *et al*., 2003; Pichoff *et al*., 2012).

To test this idea, we introduced pDSW210-FtsA-msfGFP^sw^ into *zipA1, ftsQ1, ftsI23 and ftsK44* thermosensitive mutant strains. We also introduced pDSW210 vectors as negative controls and pDSW210 derivatives (including pWM2060, pDSW210 deleted for the *gfp* gene) expressing complementing ZipA, FtsQ, FtsI, and the N terminus of FtsK (FtsK_1-220_), respectively, as positive controls. In addition, we tested *ftsK44, ftsQ1*, and *ftsI23* strains that carried pDSW210-FtsA*, as it is known that FtsA* can suppress the thermosensitivity of these alleles in addition to its prototypical ability to bypass ZipA (Schoenemann *et al*., 2018).

We were surprised to find that FtsA-msfGFP^sw^ is indeed a hypermorph, although not as strong a hypermorph as FtsA* itself. In spot viability assays, FtsA-msfGFP^sw^ permitted colony formation at the nonpermissive temperature of 42°C for *zipA1, ftsK44*, and *ftsI23* (Fig. 4). This behavior is similar to that of FtsA*, except that FtsA* could suppress all of these alleles at 42°C without IPTG induction, whereas the fusion protein required at least some level of IPTG induction except in the *ftsI23* mutant (Fig. 4). It should be noted that even ZipA and FtsI required some level of IPTG to complement their cognate thermosensitive alleles from pDSW210, although those requirements may stem from other factors such as translation efficiency (Fig. 4).

**Fig. 4.**
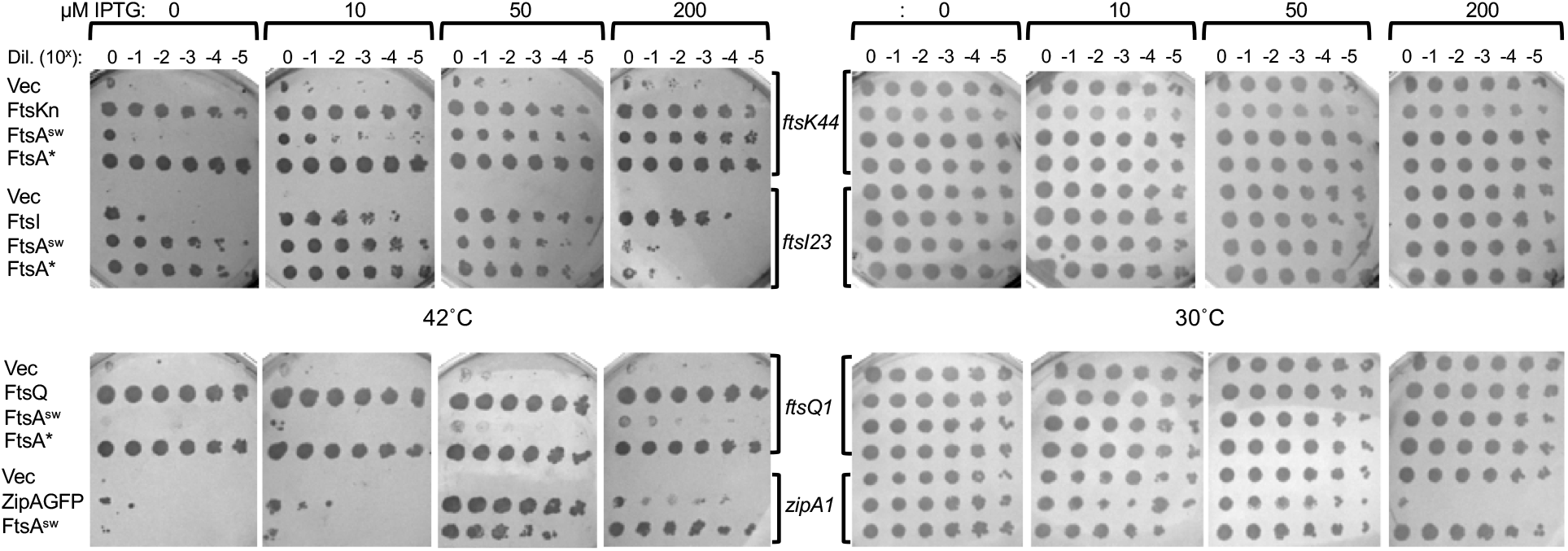
FtsA sandwich fusions can suppress thermosensitive alleles in *zipA, ftsK*, and *ftsI* but not *ftsQ*. Cultures in mid-logarithmic growth were serially diluted and spotted onto LB agar plates containing ampicillin and various concentrations of IPTG, and incubated either at 30°C or 42°C overnight. For each row: Vec is pDSW210 vector; FtsKn is pDSW210-FtsK_1-220_; FtsI is pDSW210-FtsI; FtsQ is pDSW210-FtsQ; ZipAGFP is pDSW210-ZipA-GFP; FtsA^sw^ is pDSW210-FtsA-msfGFP^sw^. Strains used, in order from top row in top section: WM6064, 5292, 6421, 3598, 5823, 5822, 6420, 6417. In order from top row in bottom section: WM5825, 5824, 6419, 6416, 6585, 6586, 6418.

As FtsA* can confer nearly normal division in cells completely devoid of ZipA (Geissler *et al*., 2003), we asked whether FtsA-msfGFP^sw^ could also completely bypass ZipA by introducing a *ΔzipA::kan* null allele into a strain expressing the sandwich fusion as the sole source of FtsA. Consistent with the requirement for IPTG to suppress *zipA1* at 42°C, kanamycin-resistant transductants were only isolated on plates with IPTG and not without IPTG (Fig. 5A). In contrast, FtsA-mCherry^sw^ could not be transduced with the *ΔzipA::kan* null allele, even with IPTG present (data not shown).

**Fig. 5.**
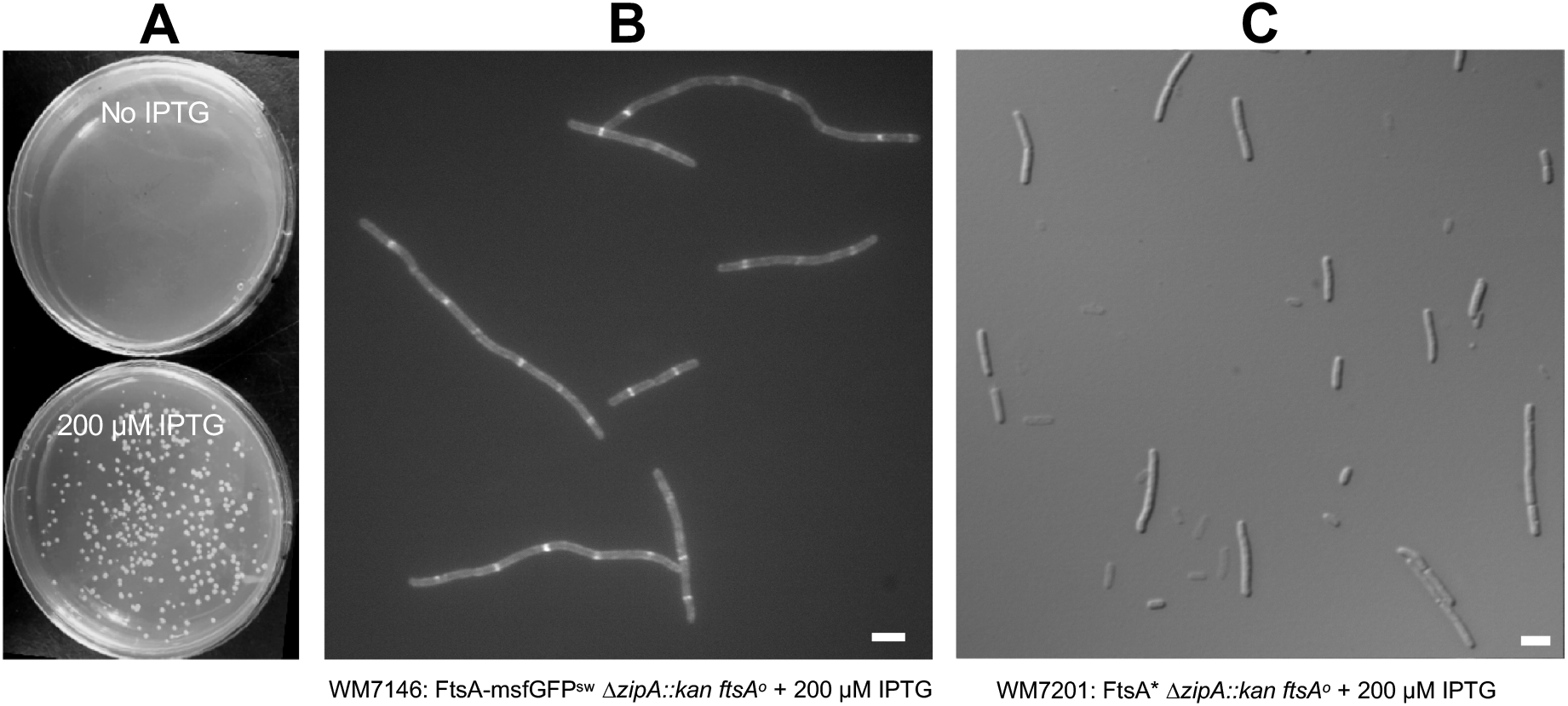
FtsA-msfGFP^sw^ can bypass a deletion of *zipA*. (A) Obtaining viable colonies at 30°C after P1-mediated transduction of the *ΔzipA::kan* allele into an *ftsA* null strain expressing FtsA-msfGFP^sw^ from pDSW210 (WM6246) requires IPTG. (B) Exponentially growing *ΔzipA::kan ftsA*^*0*^ cells expressing FtsA-msfGFP^sw^ from pDSW210 (WM7146) were grown in the presence of 200 μM IPTG, immobilized on agar pads, and subjected to fluorescence microscopy. (C) For comparison, exponentially growing *ΔzipA::kan ftsA*^*0*^ cells expressing FtsA* from pDSW210F (WM7201) were visualized with brightfield/DIC. Scale bars, 4 μm.

We then asked how efficiently FtsA-msfGFP^sw^ could divide *E. coli* cells in the absence of ZipA. Even at high IPTG induction levels, FtsA-msfGFP^sw^ was unable to restore normal division to cells with the *ΔzipA::kan* allele. Cells grown with 200 μM IPTG were a mixture of normal cells and moderately long filaments (Fig. 5B), indicating that cell division was moderately impaired but not blocked, consistent with the colony viabilities. FtsA localized to fluorescent bands at midcell in shorter cells and at multiple potential division sites in longer cells, indicating that a downstream step in divisome activity was delayed and/or impaired in the absence of ZipA. In contrast, Δ*zipA::kan ftsA*^*o*^ mutant cells expressing the *ftsA*\* allele were only slightly longer than normal (Fig. 5C), as reported previously (Geissler *et al*., 2003). This result further supports the idea that the FtsA sandwich fusions are weak hypermorphs compared with FtsA*.

We found a few interesting anomalies in the suppression assays. The first is that in contrast to FtsA*, the FtsA-msfGFP^sw^ fusion failed to suppress *ftsQ1* even with IPTG induction (Fig. 4). This was not due to protein expression problems, as expression at 42°C was still 3-4 fold higher than complementing levels of the fusion protein in the *ftsA*^*o*^ strain (Fig. 3). To validate this result, we tested the ability of the FtsA-mCherry^sw^ fusion to suppress. Like FtsA-msfGFP^sw^, FtsA-mCherry^sw^ suppressed the thermosensitivity of *ftsK44* and *zipA1* but also was unable to suppress *ftsQ1* (Fig. S1). The *ftsQ1* mutation encodes a E125K residue change, which maps to the region between the two domains of FtsQ and results in significant depletion of the protein at 42°C (Aarsman *et al*., 2005). The ability of FtsA* but not the fusion proteins to suppress *ftsQ1* suggests that FtsA* can still activate and recruit septum synthesis enzymes when FtsQ levels are very low, whereas FtsA-msfGFP^sw^ cannot.

The second interesting observation from the viability experiments is that FtsA* and FtsA-msfGFP^sw^, normally not toxic at 200 μM IPTG or even higher, became toxic at this induction level in the *ftsI23* mutant strain at 42°C (but not at 30°C) (Fig. 4). The reason for this is not clear, although it is possible that when FtsI’s septal transpeptidase activity is significantly compromised or slowed down, any increased level of divisome activation signals or recruitment of later divisome proteins negates the suppressing effects of that activation or recruitment. In particular, FtsA* may compete with FtsI for interaction with FtsN when FtsA* is at high levels in a strain with a weak FtsI. In support of this idea, FtsA* seems to interact with FtsN more strongly than does WT FtsA (Pichoff *et al*., 2018). Moreover, plasmids expressing *ftsI* that harbor mutations in its FtsN-binding domain in the periplasm could not be introduced into *ftsI23* mutant cells even at 30°C (Wissel and Weiss, 2004), indicating that inhibiting FtsI-FtsN interactions can have dominant negative effects in the presence of a weak FtsI.

### FtsA sandwich fusions persist at midcell throughout most of the septation process and rapidly relocalize to new division sites

A previous study of FtsA-GFP dynamics during the *E. coli* cell cycle suggested that FtsA persists at the septum for >85% of the division cycle but delocalized from the septum prior to cell separation (Soderstrom *et al*., 2016). As that study used a merodiploid strain expressing a C-terminally-tagged FtsA-GFP fusion that was not fully functional, we wished to test whether this also occurred with our fully functional FtsA-msfGFP^sw^ or FtsA-mCherry^sw^ as the sole FtsA in the cell. FtsA-mCherry^sw^ was more prone to photobleaching than FtsA-msfGFP^sw^, but in time-lapse series at 3 min intervals at 30°C, it is clear that FtsA-mCherry^sw^ persists at septal constrictions within 3 min of cell separation and rapidly relocalizes to midcell sites of the daughter cells (Fig. 6A-B, movies S1-S2).

**Fig. 6.**
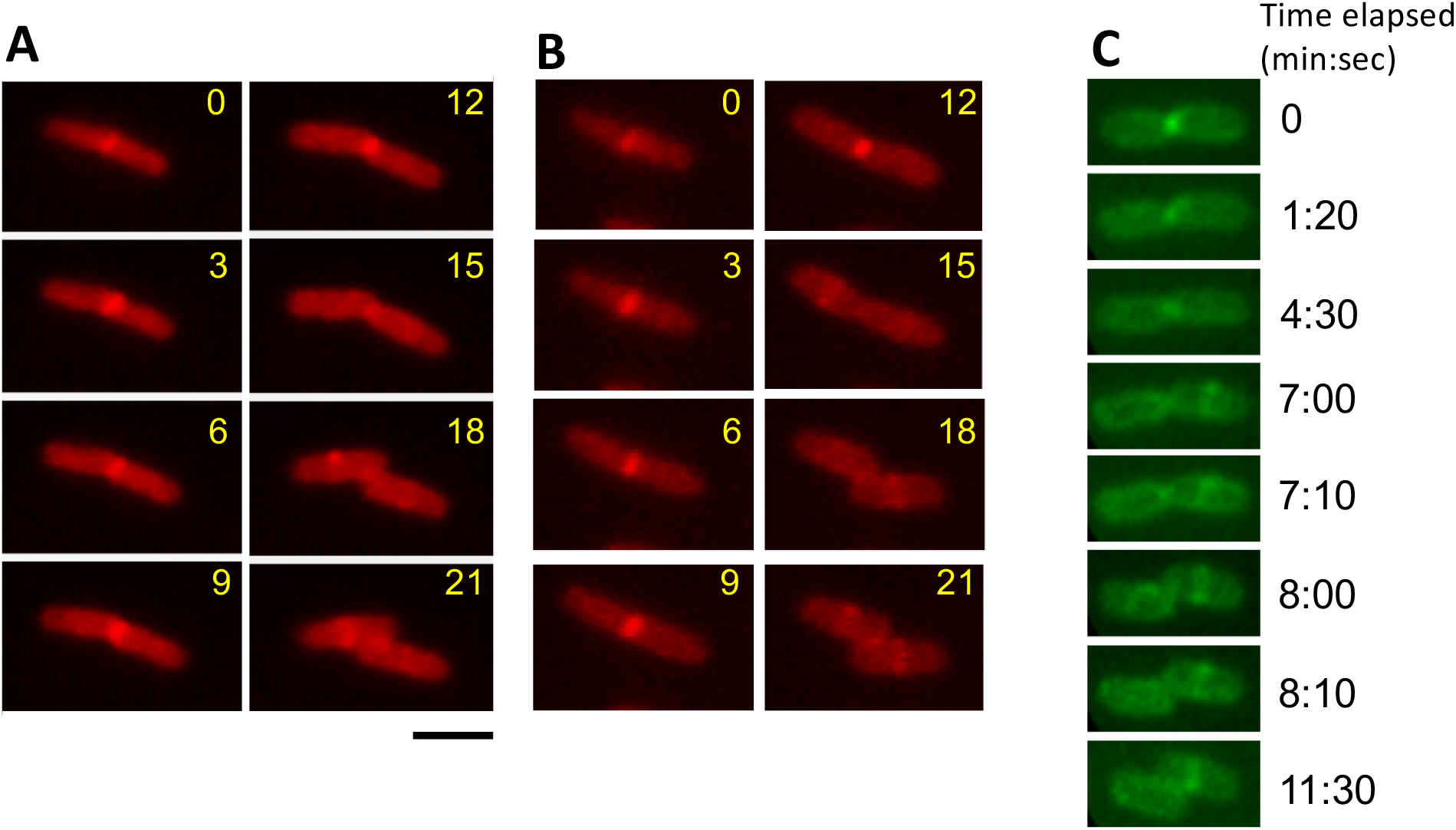
Dynamics of fluorescent FtsA during the division cycle. (A-B) Time courses of single WM4601 cells growing at ∼50-60 min doubling time imaged every 3 min, showing that FtsA-mCherry^sw^ was localized mainly at the constricting septum except for the last ∼10% (∼6 min) of the cell division cycle, where cell splitting becomes visible at 18 min in both cases. (C) Selected time points from a time course of a WM6246 cell at a late stage of septation imaged every 10 sec, showing that FtsA-msfGFP^sw^ was localized mainly at the constricting septum up to the last 3-4 min prior to visible cell separation (seen at 8:00 min). Scale bar, 2 μm.

For the FtsA-msfGFP^sw^ fusion, there was less background fluorescence, possibly due to lower proteolytic cleavage at the fluorescent protein-FtsA insert junction (data not shown). As a result, we were able to capture images at 10 sec intervals to obtain a finer temporal resolution during growth on agar pads (Fig. 6C, movie S3). The conclusions are similar to those above for the mCherry fusion: FtsA persists at the closing septum with an interval of several minutes prior to complete cell separation as judged by movement of the daughter cells, followed by immediate relocalization to midcell sites of most daughter cells. This timing is in general agreement with previous observations wherein tagged FtsA, ZipA and FtsZ persisted at the closing septum until the final ∼10% of the cell cycle, which in the case of the cells analyzed here would be ∼5 minutes.

### Detection of FtsA circumferential mobility within the Z ring

To investigate the mobility of FtsA proteins at higher spatial resolution and in greater detail, we examined the dynamic behavior of FtsA-msfGFP^sw^ in WM6246 at the cytoplasmic membrane in *E. coli* cells using total internal reflection fluorescence (TIRF) microscopy. Although photobleaching by the laser illumination precluded taking long time courses sufficient to track cell growth, we were able to capture time courses of several minutes with intervals of 2 seconds between images, allowing us to record rapid FtsA protein mobility at or near the cytoplasmic membrane in a field of multiple live cells.

One such field is shown in Fig. 7A. As expected, nearly all cells were of normal size and displayed sharp fluorescent bands or foci at midcell, indicating that TIRF could be used to capture the dynamics of functional FtsA. Strikingly, we observed two different types of dynamics. The first was outside the midcell ring: even with only 2 sec intervals between exposures, this population of FtsA moved too fast to accurately track, although it is clear that the foci formed and disassembled/moved extremely rapidly (Fig. 7B orange and red arrows, movies S4-S5). This motion also occurred in the small fraction of cells lacking clear midcell rings of FtsA-msfGFP^sw^, and indeed was observed in all the cells in this field and others (movie S4 and S6).

**Fig. 7.**
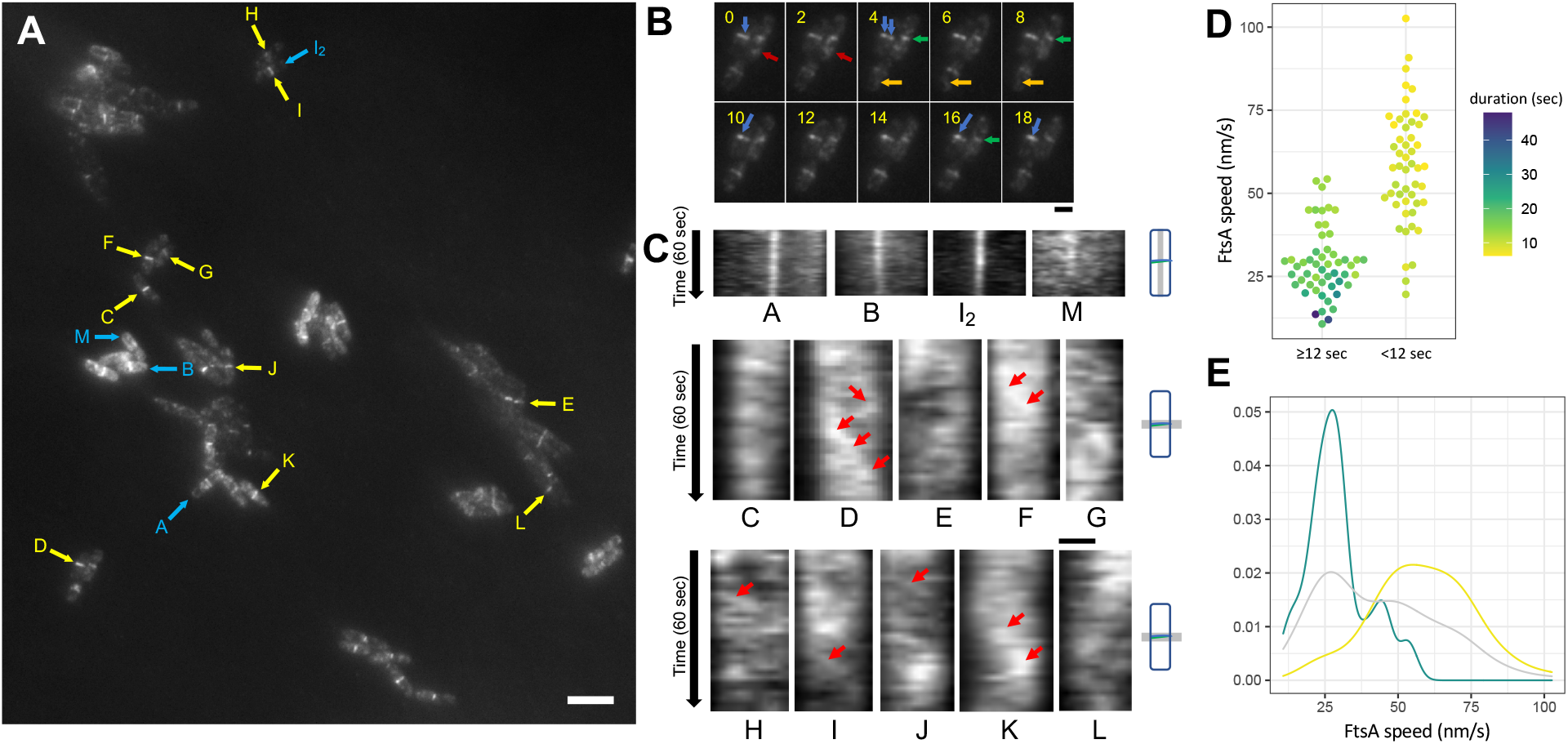
Dynamics of FtsA-msfGFP^sw^ as the sole FtsA at the membrane of *E. coli* cells observed by TIRF imaging. (A) TIRF micrograph of a field of WM6246 cells, the first frame of a 60 second time series (2 sec intervals) shown in Supplemental Movie S4. Blue arrows and letters denote cells in the time series whose fluorescence intensities are measured across their long axis, as shown in panel C (kymographs A, B, I_2_, and L). Yellow arrows and letters denote cells whose fluorescence intensities are measured across their midcell FtsA rings, also as shown in panel C (kymographs C-K, M). Scale bar, 4 μm. (B) Detail of three cells from the time course in panel A (one of which is cell D) showing the slow movement of FtsA within the midcell ring (blue and green arrows), and the fast movement of FtsA outside the ring (orange and red arrows). Time intervals are in seconds. Scale bar, 1 μm. (C) Kymographs of selected cells highlighted in panel A over the 60 second time course (2 second intervals, for a total of 31 time points along the vertical axis). The top four kymographs are from measurement along the long axis of the cell denoted by the cartoon at the right, highlighting that the bulk FtsA localization stays stable at midcell in most cells, although there is some periodicity that can be detected. The midcell band in “L” shows less stability. The bottom kymographs are from measurement across the diameter of the FtsA ring, denoted by the cartoons at the right, showing circumferential movement of bulk FtsA during the time course. Red arrows highlight circumferential motion. Scale bar, 0.42 μm. (D) Speed distribution of fluorescent FtsA-msfGFP^sw^ foci around the septum. Shown is a scatter plot with the measured velocity and lifetime of each fluorescent focus. Lifetimes are binned for <12 sec and >12 sec to illustrate the strong tendency of foci with longer lifetimes to have lower velocities that approximate those of treadmilling FtsZ. (E) The same data are represented as a kernel density estimate plot to illustrate the separation between slower moving foci with longer lifetimes and faster moving foci with shorter lifetimes: blue curve: >12 sec lifetimes; yellow curve: <12 sec lifetimes; gray curve: combined data.

The second type of dynamics occurred within the Z ring. Bulk FtsA in medial rings (A rings) was spatially stable in most cells, as depicted by the kymographs taken from the cell long axis perpendicular to the ring (Fig. 7C, top row). However, mobility of fluorescent foci around the circumference of the A ring was also apparent, as highlighted by blue and green arrows (Fig. 7B) [and in kymographs taken from thin slices across the A ring perpendicular to the cell long axis (Fig. 7C, middle and bottom rows). The high background fluorescence of the highly mobile membrane patches tended to overwhelm the fluorescence signals from the A rings, but the telltale diagonal patterns of fluorescence in parts of the kymographs suggest processive movement of FtsA around the cell circumference (Fig. 7C, yellow arrows). These diagonal patterns are similar to those of FtsZ in *E. coli* and *B. subtilis* (Bisson-Filho *et al*., 2017; Yang *et al*., 2017), although the lower signal to background for *E. coli* FtsA-msfGFP^sw^ made interpreting these much more challenging.

To determine in greater detail whether FtsA movement around the Z ring was similar to that of FtsZ, we tracked individual FtsA-msfGFP^sw^ foci over time. We found that most foci with short lifetimes within the visible TIRF field (less than 7 time points, or < 12 sec) moved rapidly, with an average speed of ∼58 ± 17 nm/sec, about double the speed previously reported for FtsZ foci (Fig. 7D). In contrast, foci with lifetimes longer than 12 sec, more likely to be part of a processively moving complex, exhibited an average speed of 29 ± 10 nm/sec and these speeds clustered in a range distinct from those of shorter-lived foci (Fig. 7D). This is within the 20-30 nm/sec range reported for treadmilling FtsZ, suggesting that at least some of the time, FtsA is moving circumferentially around midcell at the same speed as FtsZ. This estimated velocity is consistent with FtsA being a key membrane anchor for treadmilling FtsZ protofilaments; it is therefore reasonable that FtsA’s circumferential motion would match FtsZ motion. However, it should be emphasized that this is not evidence for treadmilling of FtsA oligomers, only that bulk FtsA is not locked into a fixed array beneath mobile FtsZ filaments.

### Conclusions and limitations of this study

We have developed a fully functioning fluorescent FtsA for use in studying the divisome of *E. coli*, the most studied model system for bacterial cell division, that will likely prove useful for the field. Nearly all cells with FtsA-msfGFP^sw^ as the sole copy of FtsA, even at various expression levels, are of normal length and have a fluorescent ring at midcell. These rings then constrict, during which there is a short period where FtsA leaves the constriction and relocalizes to future division sites. Using TIRF microscopy, we observed patches of FtsA-msfGFP^sw^ at midcell move relatively slowly around the septal ring, equivalent to the reported speed of FtsZ treadmilling polymers (Yang *et al*., 2017). In addition, we discovered that FtsA-msfGFP^sw^ assembles into multiple foci outside the ring that move much more rapidly around the cell periphery, with estimated turnover times <2 sec and possibly much lower than that. The structure and potential function of these dynamic peripheral complexes are unknown, but they may be related to the transient non-ring complexes of FtsZ that contain significant quantities of membrane associated FtsZ (Walker *et al*., 2020; Margolin, 2020).

Although the sandwich fusion protein described here will undoubtedly be useful for additional studies, it has some limitations. The non-ring localization of FtsA-msfGFP^sw^, while intriguing, increases the background fluorescence surrounding the A ring, posing challenges for single molecule localization and tracking of FtsA in the A ring. The rapid motion of this non-ring FtsA makes it difficult to track mobile FtsA-msfGFP^sw^ foci, particularly as they migrate away from the cell surface and thus out of the ∼200 nm deep TIRF detection zone, and to distinguish these foci from others closer to each other than the resolution limit. Potential ways to overcome this are to use other fluorophores that have higher efficiencies while retaining the activity of the FtsA protein, such that the fluorescent FtsA can be expressed at lower levels closer to those of the native protein.

Other continuing challenges include fusions to fluorescent proteins in general. While they have been invaluable for tracking protein localization in living cells and can often replace the untagged proteins in vivo, they often are not fully functional. This is a particular issue for proteins such as FtsZ and FtsA that form highly regulated polymeric structures. The original C-terminally tagged FtsZ-GFP and FtsA-GFP could not replace the native proteins (Ma *et al*., 1996), although they could be used to visualize FtsZ and FtsA localization in merodiploids. The development of a FtsZ-mNeonGreen sandwich fusion that can substitute for the native FtsZ was an important advance, but this fusion still does not divide cells as efficiently as native FtsZ, with detectable defects in cell length and morphology when grown in rich medium (Moore *et al*., 2017). These defects are suppressed at slower growth rates in minimal medium, but are not absent.

In contrast, our FtsA sandwich fusions seem to have few if any division defects even during rapid growth in LB or when overproduced. Nonetheless, they are not completely WT in function, as they exhibit gain-of-function properties. This should not be too surprising, as defects in normal assembly of FtsA oligomers caused by steric interference of the inserted fluorescent protein might result in a paradoxical gain of function. FtsA in the minirings is probably oriented with its 1A subdomain on the outside (Nierhaus *et al*., 2022), and the fluorescent protein insertion is in the IIB subdomain on the opposite side of the FtsA molecule. As a result, we speculate that the bulky beta barrel structure of the fluorescent protein would be oriented toward the lumen of any putative miniring structure *in vivo*, and the need to pack 12 such structures in the limited space of the lumen might prevent formation of a closed miniring, but not curved or straight oligomers with free ends. Such behavior might shift the population of FtsA sandwich fusions towards curved oligomers or double stranded filaments, either of which seem to be an activated form of FtsA (Schoenemann *et al*., 2018; Nierhaus *et al*., 2022). The inability of the sandwich fusions to suppress *ftsQ1* indicates that although the fusions can divide cells normally, they still require functions of the divisome checkpoint that are normally bypassed by FtsA*. Future studies will be needed to determine the exact structural basis of these gain of function properties.

*B. subtilis* FtsA was shown to treadmill alongside FtsZ *in vivo* (Bisson-Filho *et al*., 2017), suggesting that *B. subtilis* FtsA assembles into oligomers with polarity. An outstanding question in the field is whether *E. coli* FtsA also treadmills. Evidence from *in vitro* structures on membrane surfaces argues against it. First, *E. coli* FtsA minirings or arcs likely do not treadmill, as treadmilling requires free ends not observed in minirings, and treadmilling at the termini of FtsA arcs would result in going around in circles and not processive movement around the septum. Second, the double stranded filaments of *E. coli* FtsA, activated either by mutation or by binding with the cytoplasmic domain of FtsN, are antiparallel, which should not be capable of treadmilling as they have no net polarity (Nierhaus *et al*., 2022). Therefore, we propose that FtsA moves around the septum by other mechanisms. Minirings and arcs may move by rapidly assembling and disassembling to keep up with treadmilling FtsZ polymers, and FtsA double stranded filaments may act as a rudder and be pushed around like MreB filaments (van den Ent *et al*., 2014; Hussain *et al*., 2018), as recently proposed (Nierhaus *et al*., 2022). The population of fluorescent FtsA foci that we measured to move at speeds similar to treadmilling FtsZ suggests that these FtsA molecules, in whatever oligomeric form, function as a moveable track for FtsZ. The faster-moving FtsA may reflect a population that is not engaged in tethering FtsZ, perhaps because of overexpression. Future investigations, using these and potentially optimized fluorescent versions of FtsA, will be necessary to gain insight into these potential structures and mechanisms.

## METHODS

### Strains and cell growth conditions

All strains and plasmids are listed in Table S1. *E. coli* strains Top10 and XL1-Blue were used as the host strains for molecular cloning. Cells were cultured in Lysogeny Broth (LB) agar or broth containing 0.5% NaCl at 30°C or 42°C except where indicated. LB agar or broth (with 0.5 % NaCl) was supplemented with ampicillin (50 μg/ml), kanamycin (25 μg/ml) tetracycline (10 μg/ml) and/or isopropylthio-beta-galactoside (IPTG) at the indicated concentrations when needed.

### Construction of FtsA sandwich fusion plasmids and strains

To engineer *ftsA*-mCherry^sw^ (pWM4450), the mCherry sequence was inserted between encoded residues S267 and I268 of *E. coli* FtsA with a 7-residue flanking sequence (encoding ELSGAPG) at the 5’ end of the insert, and the 8-residue SSGAPGGS sequence at the 3’ end of the insert, by InFusion cloning (Takara Bio). Primers 1965 and 1966 were used as the inner primers and 1967 and 1968 as the outer primers for amplifying the *ftsA* gene from pWM2785; all primers are listed in Table S2. The resulting plasmid, a derivative of pWM2060 which is a GFP-less version of pDSW210, contains the mCherry gene flanked by SacI and BamHI sites. To replace the mCherry insert with msfGFP, the msfGFP gene from pNOMreB-msfGFP^sw^ (WM5899) was PCR-amplified using primers 2156 and 2157, cleaved with SacI and BamHI, and cloned into pWM4450 cleaved with the same enzymes. The resulting plasmid, pWM6246, contained the 6-residue linkers ELSGSS at the N-terminal junction between msfGFP and FtsA and SGAPGS at the C-terminal junction.

To introduce the *ftsA*^*o*^ null allele into strains with plasmids expressing *ftsA* fusions, a phage P1 lysate carrying *ftsA*^*o*^, which carries a frameshift mutation resulting from a fill-in and religation at the unique Bgl II site in *ftsA*, was used to cotransduce a *leuO*::Tn*10* marker that maps ∼20 kb away. Tc^R^ transductants were screened for the loss of the Bgl II site by colony PCR, using primers specific for the regions flanking the chromosomal *ftsA* locus. The frequency of cotransduction was ∼50%.

To transfer the msfGFP^sw^ fusion to the lower expression plasmid pSEB440, plasmid pWM6246 was used as a template with primers 2580 and 2581 to amplify a fragment with msfGFP and flanking FtsA sequence containing native 5’ KpnI and 3’ AscI sites. This fragment was digested with KpnI and AscI, then cloned into pSEB440 cut with the same enzymes, selected for chloramphenicol resistance, and screened for the ability to form fluorescent midcell bands upon addition of IPTG. This strain was saved as WM7118.

To introduce the *ftsA*^*o*^ or *ftsA*^*o*^ + *ftsL*\* alleles into WM7118, P1 lysates from either WM4952 or WM4953 that contain a *leuO*::Tn*10* marker linked to *ftsA*^*o*^ (WM4952) or both *ftsL*\* and *ftsA*^*o*^ (WM4953) were used to transduce WM6776 (pSEB440 positive control) or WM7118, selecting for Cm^R^ and Tc^R^ in the presence of 500 μM IPTG. Transductant colonies for all four transductions were then purified on plates with chloramphenicol, tetracycline and 500 μM IPTG and screened on plates lacking IPTG.

Of the WM4952 transductants in WM6776, 4 out of 8 grew on no IPTG, compared with 3 out of 8 for WM7118, consistent with the ∼50% linkage between the Tc^R^ Tn10 marker and *ftsA*^*o*^ and the requirement of IPTG for expression of FtsA or FtsA-msfGFP^sw^ from this plasmid. In contrast, 8 out of 8 WM6776 transductants grew on 500 μM IPTG, compared with 3 out of 8 WM7118 transductants, indicating that strong induction with IPTG was insufficient to allow FtsA-msfGFP^sw^ to function as the sole FtsA when expressed from pSEB440.

Of the WM4953 transductants in WM6776, 5 out of 8 grew well without IPTG, and the other 3 grew to small colonies, consistent with the ability of *ftsL*\* to allow some survival of plasmid *ftsA*-expressing cells in the absence of IPTG (*ftsL*\* is located between the Tn10 and *ftsA* and thus will always be present in Tc^R^ transductants carrying *ftsA*^*o*^). In contrast, 4 out of 8 WM7118 transductants grew well without IPTG, while the other 4 had very low viability, indicating that even the presence of *ftsL*\* did not allow viability of the strain expressing FtsA-msfGFP^sw^ from pSEB440 without IPTG. All transductants of WM6776 and WM7118 grew robustly with 500 μM IPTG, indicating that the addition of *ftsL*\* allowed the pSEB440 expressing FtsA-msfGFP^sw^ cells to divide efficiently in the absence of native *ftsA*. PCR for the presence of absence of the Bgl II site within the *ftsA* gene confirmed the that the *ftsA*^*o*^ allele was acquired in the strains.

To test whether the sandwich fusions could bypass the complete loss of *zipA*, we used a P1 lysate from WM1657, carrying a *ΔzipA::kan* allele along with its required *ftsA*\* suppressor, to transduce WM4550 and WM6246. Transductants were selected on LB plates with kanamycin + 200 μM IPTG. WM6246 yielded transductants at the expected frequency, whereas WM4550 yielded no viable transductants. To construct the positive control strain WM7201 carrying *ΔzipA::kan* plus *ftsA*\* expressed from a pDSW210 derivative, WM5732 was transduced with the *ΔzipA::kan* allele. WM5731, expressing WT *ftsA* from the same plasmid backbone, was transduced with *ΔzipA::kan* in parallel but yielded no transductants as expected.

### Other plasmid constructions

The FtsK_1-200_-FLAG plasmid pWM5292 for complementation of *ftsK44* was constructed by amplifying the *E. coli ftsK* gene with primers FtsKnFlagF and FtsKnFlagR followed by cloning the PCR product into pDSW210 between the SacI and XbaI sites. The resulting FtsK_1-200_-FLAG is under IPTG control and has a stop codon after the FLAG peptide.

### Serial dilution and plating assays

Overnight cell cultures were diluted 1:200 and grown to OD_600_ = 0.2-0.4. Cell densities were normalized between strains in 200 μL before being serially diluted 10-fold and spotted on agar plates either with a 0.4 μL-hanging drop 48-prong cell replicator or a multichannel pipettor (4 μl). All agar plates contained LB supplemented with the appropriate antibiotics and different concentrations of IPTG.

### Immunoblot analysis

Overnight cultures were diluted 1:200 in 4 mL LB and grown for 2.5 h at 30°C. Cultures with 42°C treatments were then split in half and placed in a 42°C shaking water bath incubator and all cultures were incubated for an additional 1h. Cultures were pelleted and resuspended in 1x SDS-PAGE sample loading buffer at a concentration of 0.01 OD_600_/μL. Samples were boiled for 10 min, then 7.5 μL of each was separated by SDS-PAGE using 12.5% tris-glycine polyacrylamide gels and Laemmli running buffer. Samples were transferred to 0.2 μm nitrocellulose membranes using a Mini Trans-Blot apparatus (BioRad) at 200 mA for 55 min in Towbin transfer buffer modified to include 0.1% SDS and 20% ethanol. Blots were stained by Ponceau to assess total protein levels, then blocked using 3% BSA in TBST and incubated with rabbit anti-FtsA (1:2500) antibodies. After washes, blots were incubated with goat anti-rabbit-HRP (1:10000), developed with Pierce ECL substrate and imaged using a ChemiDoc MP system.

### Microscopy and image analysis

For standard epifluorescence imaging, 3 μl of cells in mid-logarithmic phase were spotted on a thin 1% agarose pad containing LB and covered with a cover glass. Fluorescence micrographs were acquired with an Olympus BX63 microscope equipped with a 100X N.A. 1.4 objective and Hamamatsu C11440 ORCA-Spark digital CMOS camera, using a FITC filter cube for GFP and a TRITC filter cube for mCherry. Images were captured with cellSens software (Olympus). For time lapse imaging with epifluorescence, 3 μl of cells in mid-logarithmic phase were spotted in the center of a culture dish fitted with a cover glass (Mattek Corp.). A thin slab of LB agar or LB agarose was then placed on top of the cells to immobilize them and provide nutrients. The culture dish was then placed onto the stage of an Olympus IX-70 inside a WeatherStation temperature-controlled incubator at 30°C. Time-lapse images of growing cells were acquired with a Hamamatsu ORCA CCD camera using GFP or mCherry filter cubes and SlideBook software from Intelligent Imaging Innovations, Inc. Further image processing was done with Fiji software (Schindelin *et al*., 2012).

For TIRF imaging, we used a Nikon n-STORM workstation at the UTHealth Houston Center for Advanced Microscopy. WM6246 cells in logarithmic growth expressing FtsA-msfGFP^sw^ were immobilized on Mattek dishes as described above, then imaged with a Nikon n-STORM Eclipse T1 inverted microscope equipped with TIRF HP APO 100X objective and a 70 mW DPSS laser line at 488 nm. Images were acquired with a Hamamatsu digital camera and analyzed with Nikon Elements software.

Further image processing and kymograph analysis were done with Fiji. Kymographs were generated for a total of 220 cells with midcell FtsA fluorescence using the “Multi Kymograph” function with a line 3 pixels thick (∼180 nm) drawn over each midcell band. To avoid measuring spurious fluorescence from unassociated FtsA foci moving throughout the entire cell membrane, the resulting kymographs were compared with the time course of each cell to identify distinct foci with consistent circumferential motion over at least 6 seconds. The lengths and angles of linear tracks in the kymographs were measured for a total of 104 such foci. Data was further analyzed using R (v4.2.1) and visualized using the ggplot2 (v3.3.6) and ggbeeswarm (v0.7.0.900) packages.

## Acknowledgements

We thank Yi Liu, Natalie R. Williams, Kara M. Schoenemann, Mwidy Mounange-Badimi, and Steven L. Distelhorst for strain constructions and generating preliminary data. We are grateful to Kyung-Tae Park and Joe Lutkenhaus for providing plasmid pSEB440 and Ben Bratton and Joshua Shaevitz for providing plasmid pNOMreB-msfGFP^sw^. We thank Travis Moore, Olga Chumakova, and the UTHealth Houston Center for Advanced Microscopy for their valuable expertise and assistance with TIRFM. This study was supported by NIH grant R35GM131705.

---

**Fig S1:**
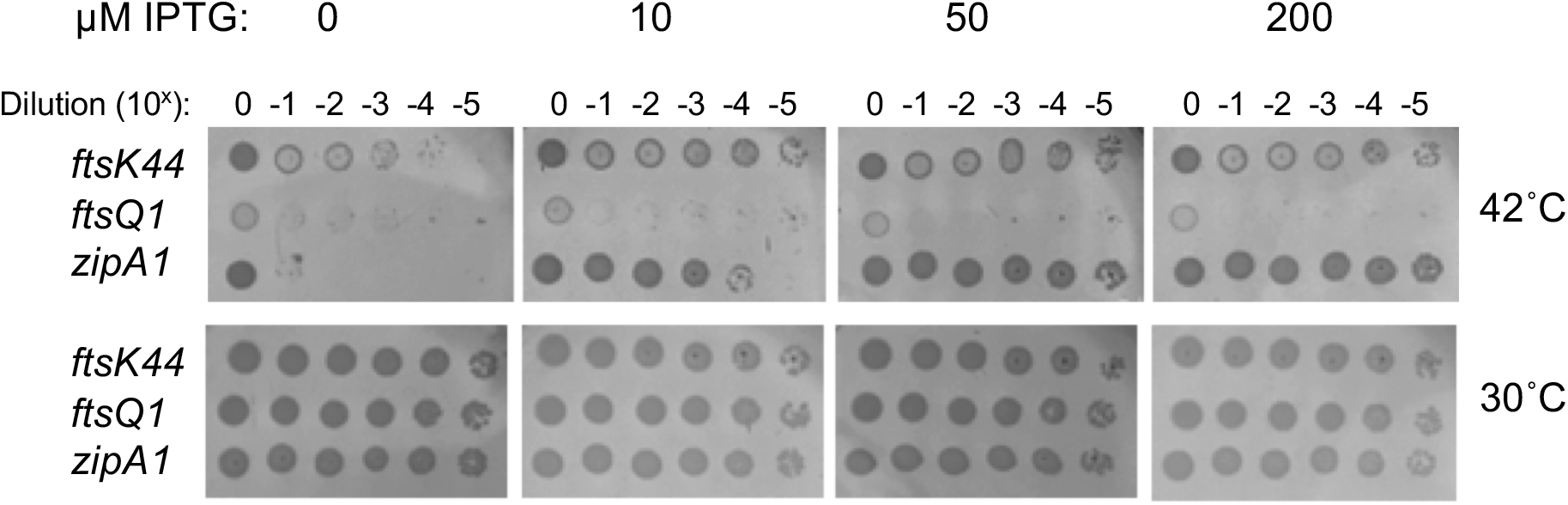
FtsA-mCherry^sw^ suppression of divisome mutants is similar to that of FtsA-msfGFP^sw^. Cells of WM7158, 7156, and 7157 (pDSW210-FtsA-mCherry^sw^ in *ftsK44, ftsQ1*, and *zipA1* thermosensitive mutant strain backgrounds, respectively) growing in mid-logarithmic phase were serially diluted and spotted on LB + Ampicillin with different IPTG concentrations and incubated for 20 h at 30°C or 42°C.

**Table S1.**
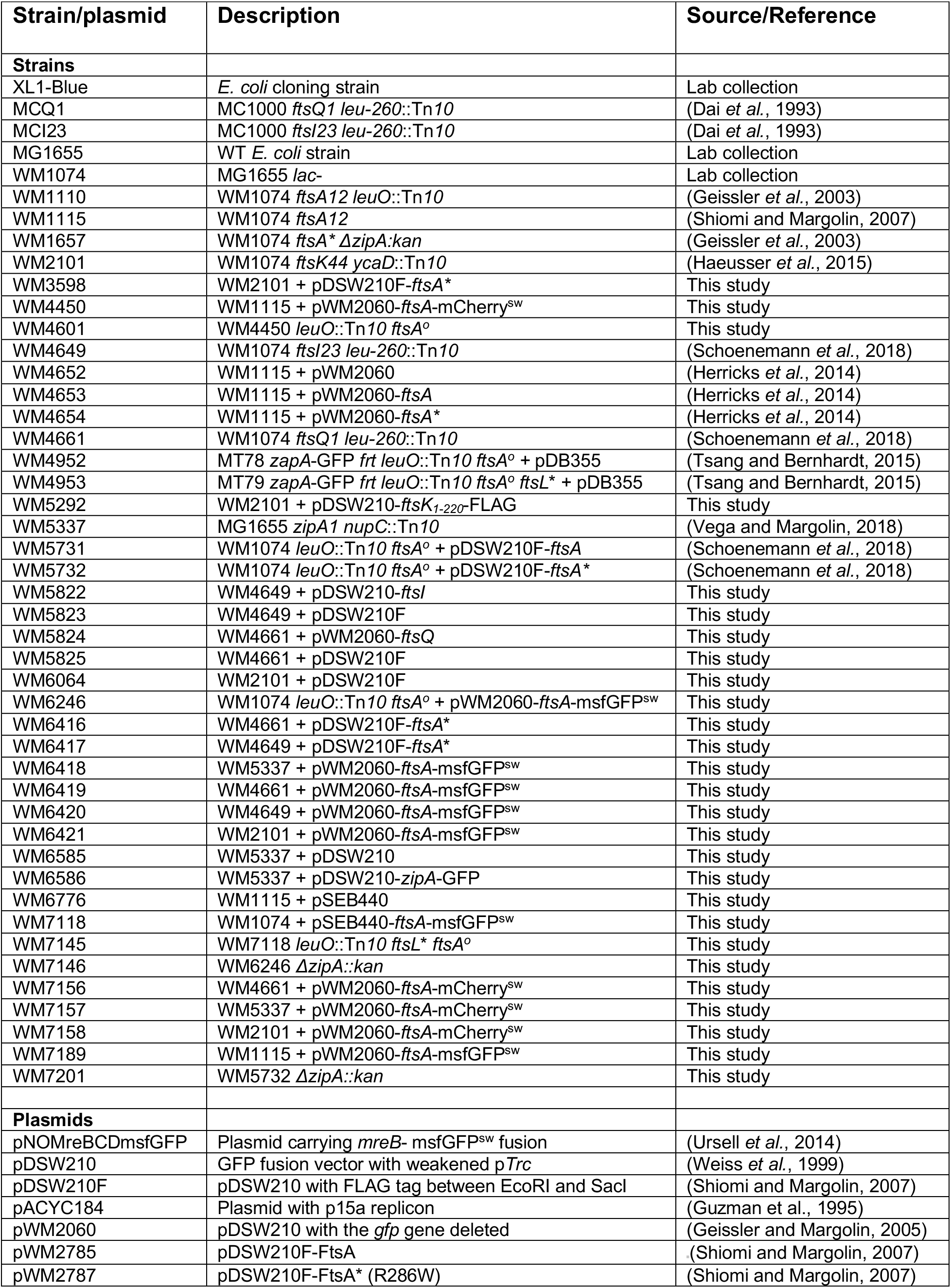
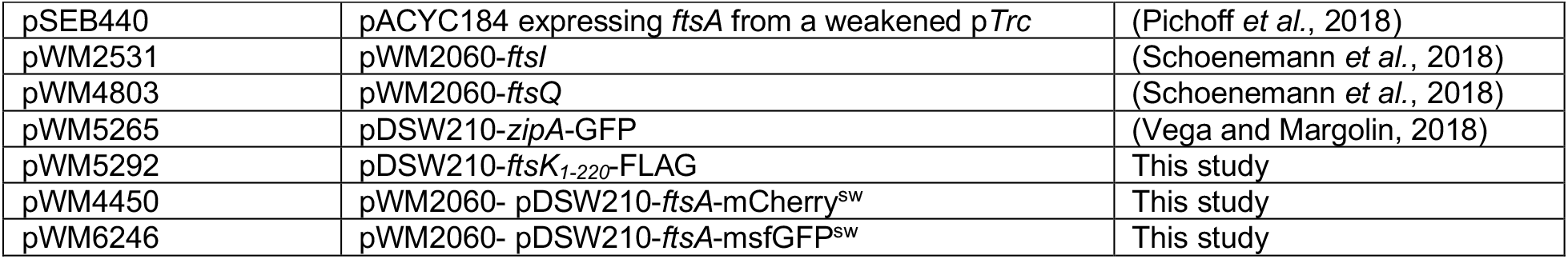
Strains and plasmids used in this study.

| Strain/plasmid | Description | Source/Reference |
| --- | --- | --- |
| **Strains** |  |  |
| XL1-Blue | *E. coli* cloning strain | Lab collection |
| MCQ1 | MC1000 *ftsQ1 leu-260::Tn10* | (Dai *et al.*, 1993) |
| MCI23 | MC1000 *ftsI23 leu-260::Tn10* | (Dai *et al.*, 1993) |
| MG1655 | WT *E. coli* strain | Lab collection |
| WM1074 | MG1655 *lac-* | Lab collection |
| WM1110 | WM1074 *ftsA12 leuO::Tn10* | (Geissler *et al.*, 2003) |
| WM1115 | WM1074 *ftsA12* | (Shiomi and Margolin, 2007) |
| WM1657 | WM1074 *ftsA\* ΔzipA::kan* | (Geissler *et al.*, 2003) |
| WM2101 | WM1074 *ftsK44 ycaD::Tn10* | (Haeusser *et al.*, 2015) |
| WM3598 | WM2101 + pDSW210F-*ftsA\** | This study |
| WM4450 | WM1115 + pWM2060-*ftsA-mCherry<sup>sw</sup>* | This study |
| WM4601 | WM4450 *leuO::Tn10 ftsA<sup>o</sup>* | This study |
| WM4649 | WM1074 *ftsI23 leu-260::Tn10* | (Schoenemann *et al.*, 2018) |
| WM4652 | WM1115 + pWM2060 | (Herricks *et al.*, 2014) |
| WM4653 | WM1115 + pWM2060-*ftsA* | (Herricks *et al.*, 2014) |
| WM4654 | WM1115 + pWM2060-*ftsA\** | (Herricks *et al.*, 2014) |
| WM4661 | WM1074 *ftsQ1 leu-260::Tn10* | (Schoenemann *et al.*, 2018) |
| WM4952 | MT78 *zapA-GFP frt leuO::Tn10 ftsA<sup>o</sup> + pDB355* | (Tsang and Bernhardt, 2015) |
| WM4953 | MT79 *zapA-GFP frt leuO::Tn10 ftsA<sup>o</sup> ftsL\* + pDB355* | (Tsang and Bernhardt, 2015) |
| WM5292 | WM2101 + pDSW210-*ftsK1-220-FLAG* | This study |
| WM5337 | MG1655 *zipA1 nupC::Tn10* | (Vega and Margolin, 2018) |
| WM5731 | WM1074 *leuO::Tn10 ftsA<sup>o</sup> + pDSW210F-ftsA* | (Schoenemann *et al.*, 2018) |
| WM5732 | WM1074 *leuO::Tn10 ftsA<sup>o</sup> + pDSW210F-ftsA\** | (Schoenemann *et al.*, 2018) |
| WM5822 | WM4649 + pDSW210-*ftsI* | This study |
| WM5823 | WM4649 + pDSW210F | This study |
| WM5824 | WM4661 + pWM2060-*ftsQ* | This study |
| WM5825 | WM4661 + pDSW210F | This study |
| WM6064 | WM2101 + pDSW210F | This study |
| WM6246 | WM1074 *leuO::Tn10 ftsA<sup>o</sup> + pWM2060-ftsA-msfGFP<sup>sw</sup>* | This study |
| WM6416 | WM4661 + pDSW210F-*ftsA\** | This study |
| WM6417 | WM4649 + pDSW210F-*ftsA\** | This study |
| WM6418 | WM5337 + pWM2060-*ftsA-msfGFP<sup>sw</sup>* | This study |
| WM6419 | WM4661 + pWM2060-*ftsA-msfGFP<sup>sw</sup>* | This study |
| WM6420 | WM4649 + pWM2060-*ftsA-msfGFP<sup>sw</sup>* | This study |
| WM6421 | WM2101 + pWM2060-*ftsA-msfGFP<sup>sw</sup>* | This study |
| WM6585 | WM5337 + pDSW210 | This study |
| WM6586 | WM5337 + pDSW210-*zipA-GFP* | This study |
| WM6776 | WM1115 + pSEB440 | This study |
| WM7118 | WM1074 + pSEB440-*ftsA-msfGFP<sup>sw</sup>* | This study |
| WM7145 | WM7118 *leuO::Tn10 ftsL\* ftsA<sup>o</sup>* | This study |
| WM7146 | WM6246 *ΔzipA::kan* | This study |
| WM7156 | WM4661 + pWM2060-*ftsA-mCherry<sup>sw</sup>* | This study |
| WM7157 | WM5337 + pWM2060-*ftsA-mCherry<sup>sw</sup>* | This study |
| WM7158 | WM2101 + pWM2060-*ftsA-mCherry<sup>sw</sup>* | This study |
| WM7189 | WM1115 + pWM2060-*ftsA-msfGFP<sup>sw</sup>* | This study |
| WM7201 | WM5732 *ΔzipA::kan* | This study |
| **Plasmids** |  |  |
| pNOMreBCDmsfGFP | Plasmid carrying *mreB- msfGFP<sup>sw</sup>* fusion | (Ursell *et al.*, 2014) |
| pDSW210 | GFP fusion vector with weakened pTrc | (Weiss *et al.*, 1999) |
| pDSW210F | pDSW210 with FLAG tag between EcoRI and SacI | (Shiomi and Margolin, 2007) |
| pACYC184 | Plasmid with p15a replicon | (Guzman *et al.*, 1995) |
| pWM2060 | pDSW210 with the *gfp* gene deleted | (Geissler and Margolin, 2005) |
| pWM2785 | pDSW210F-*FtsA* | (Shiomi and Margolin, 2007) |
| pWM2787 | pDSW210F-*FtsA\** (R286W) | (Shiomi and Margolin, 2007) |
| pSEB440 | pACYC184 expressing *ftsA* from a weakened pTrc | (Pichoff *et al.*, 2018) |
| pWM2531 | pWM2060-*ftsI* | (Schoenemann *et al.*, 2018) |
| pWM4803 | pWM2060-*ftsQ* | (Schoenemann *et al.*, 2018) |
| pWM5265 | pDSW210-*zipA*-GFP | (Vega and Margolin, 2018) |
| pWM5292 | pDSW210-*ftsK*<sub>1-220</sub>-FLAG | This study |
| pWM4450 | pWM2060- pDSW210-*ftsA*-mCherry<sup>SW</sup> | This study |
| pWM6246 | pWM2060- pDSW210-*ftsA*-msfGFP<sup>SW</sup> | This study |

**Table S2.**
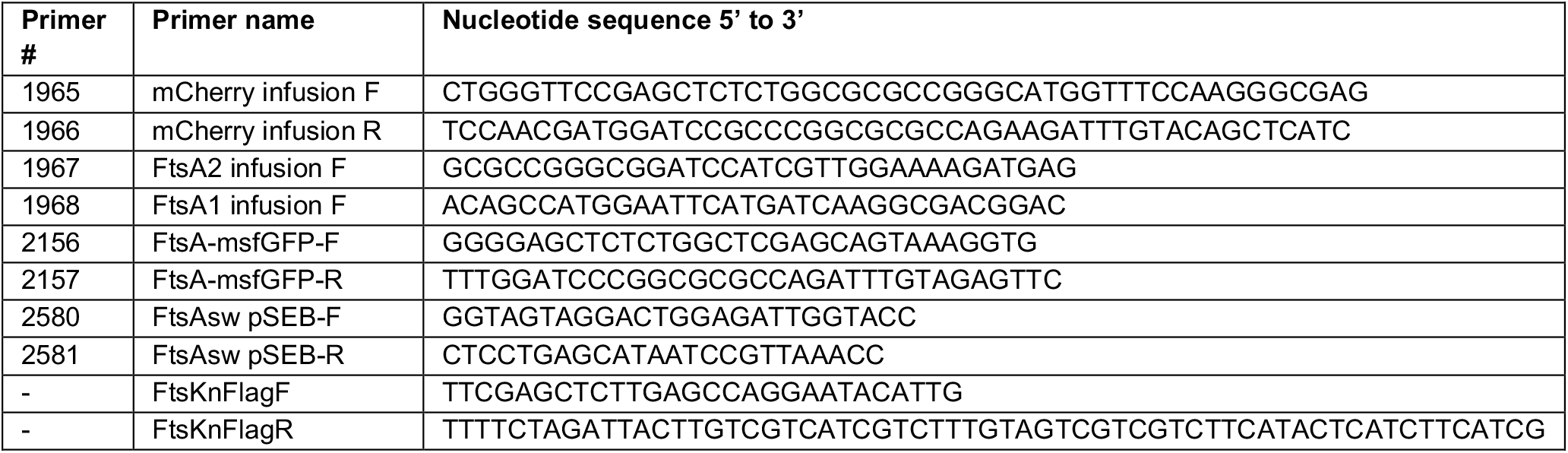
Primers used in this study.

